# Morphological fingerprints enable machine learning–based inference of neuroblastoma cell states without transcriptomics

**DOI:** 10.64898/2026.05.12.724731

**Authors:** Vic Zamloot, Yuanzhong Pan, JinSeok Park

## Abstract

Inference of cancer cell states is essential for understanding oncogenic mechanisms and predicting clinical outcomes, yet current reliance on transcriptomic profiling limits scalability and real-time monitoring. Here, we show that cell morphology provides a low-dimensional, observable representation of cellular identity and its dynamics. Using neuroblastoma (NB) as a model system, we establish a machine learning- morphology profiling framework that infers adrenergic (ADRN) and mesenchymal (MES) cell states directly from high-dimensional morphological fingerprints without reliance on transcriptomic measurements. By benchmarking against single-cell RNA sequencing (scRNA-seq), we demonstrate that morphology-defined states closely align with transcriptomic profiles at single-cell resolution. We further show that cell state transitions are represented as continuous trajectories within a morphology-defined state space. Perturbations targeting distinct regulatory layers, including ROCK signaling and epigenetic regulation via EZH2, drive convergent trajectories along a shared phenotypic axis. Together, these results establish cell morphology as a scalable and non-destructive readout of cell state with machine learning providing a unified framework for high-throughput phenotyping and real-time tracking of cancer cell state plasticity.

## Introduction

Intra- and inter-tumoral heterogeneity remains a fundamental challenge in cancer biology, complicating the understanding of oncogenic mechanisms and the prediction of clinical outcomes ^1-3^. While single-cell RNA sequencing (scRNA-seq) has enabled high-resolution identification of cellular subpopulations ^4-6^, its widespread application is limited by cost, destructive sampling, and the inability to capture real-time, longitudinal dynamics. These limitations underscore the need for scalable and non-invasive approaches to monitor cellular states and their plasticity.

Cell morphology reflects the integrated output of oncogenic signaling and cytoskeletal organization ^7-10^. However, morphological phenotypes are inherently high-dimensional and complex, making it challenging to directly interpret their relationship to underlying cell states. We therefore hypothesized that morphology defines a low-dimensional state space that encodes cellular identity and its dynamics. To test this, we developed a machine-learning based framework that extracts latent information from multidimensional “morphological fingerprints” and enables quantitative mapping of cell states.

Neuroblastoma (NB), a pediatric malignancy accounting for ~15% of childhood cancer deaths, provides an ideal model system to investigate cell state plasticity, as tumor heterogeneity is largely driven by non-genetic state transition rather than mutational burden^11^. NB cells interconvert between two distinct states: a lineage-committed adrenergic (ADRN) state and an undifferentiated mesenchymal (MES) state associated with poor prognosis^12,13^. Accurate identification and tracking of these states are therefore critical for understanding NB progression and therapeutic response.

Here, we establish a morphology-based machine learning framework to infer NB cell states directly from high-dimensional morphological fingerprints without reliance on transcriptomic measurements. By linking morphological phenotypes to transcriptomic states, we show that cell identity can be represented as a low-dimensional morphology-defined state space and that state transitions occur as continuous trajectories within this space at single-cell resolution. We further demonstrate that perturbations at the signaling and epigenetic level, including ROCK and EZH2 inhibition, drive convergent trajectories indicating the plasticity. Together, these findings establish cell morphology as a scalable and non-destructive readout of cell state with machine learning, enabling high-throughput phenotyping and real-time monitoring of cancer cell state plasticity.

## Results

### ScRNA-seq analysis reveals ADRN and MES states in NB cell lines

To characterize the NB heterogeneity comprising MES and ADRN states, we analyzed publicly available scRNA-seq data from multiple NB cell lines (GSE229224)^14^. Principal component analysis (PCA) of the integrated data set revealed that cells primarily cluster by cell line identity, while also exhibiting continuous transcriptional variation within each line (**Fig. 1A**). To further investigate MES and ADRN states, we quantified ADRN and MES scores of each cell using single-sample gene set enrichment analysis (ssGSEA)^15,16^ with reference gene sets of ADRN and MES signatures previously defined by van Groningen *et al*.^13^ Projection of these scores onto the PCA space demonstrated opposing gradients, consistent with their distinct molecular features (**Fig. 1B; Supplementary Data 1**).

**Figure 1.**
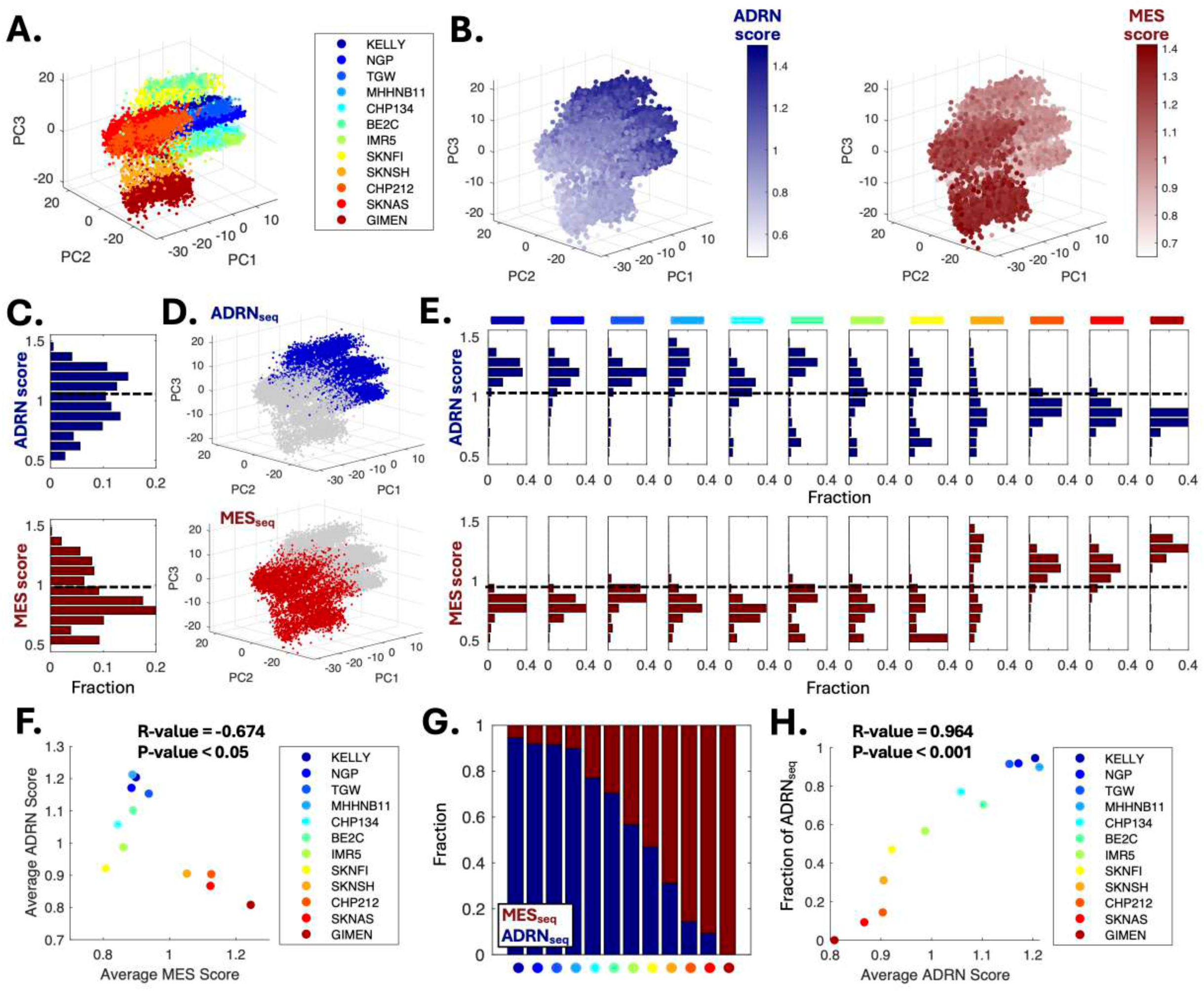
Single-cell sequencing analyses of neuroblastoma (NB) cell lines identify distinct ADRN and MES profiles. **(A)** Three-dimensional principal component analysis (PCA) of publicly available single-cell RNA sequencing (scRNA-seq) data from NB cell lines. Each color represents a corresponding cell line. Data are from GSE229224. **(B)** PCA plots showing ADRN (left) and MES (right) scores calculated using single-sample Gene Set Enrichment Analysis (ssGSEA). **(C)** Histograms of ADRN (top) and MES (bottom) scores for individual cells. Dashed lines indicate thresholds separating high- and low-score groups. Thresholds were determined using a machine learning–based algorithm (k-means, k = 2). **(D)** PCA plots highlighting an ADRN subgroup defined by high ADRN scores (ADRN_seq_, top) and an MES subgroup defined by low ADRN scores (MES_seq_, bottom) in the scRNA-seq analysis. **(E)** Histograms of ADRN (top) and MES (bottom) scores for individual cells in each cell line. Color bars indicate the corresponding cell lines shown in Panel (A). Dashed lines represent the thresholds defined in Panel C. **(F)** Scatter plot showing the relationship between average ADRN and MES scores across cell lines. **(G)** Fraction of ADRN_seq_ and MES_seq_ cells defined by ADRN scores in each cell line. Each colored dot corresponds to a cell line shown in Panel (F). **(H)** Scatter plot showing the relationship between average ADRN score and the fraction of ADRN_seq_ cells in each cell line. In Panel (F) and (H), R and P values were calculated using Pearson’s correlation coefficient.

At the single-cell level, ADRN and MES scores displayed broad distributions; notably, ADRN scores exhibited a distinct bimodal pattern, suggesting the existence of two discrete subpopulations (**Fig. 1C**). Using a data-driven thresholding approach via k-means clustering (k = 2) ^17-19^ of ADRN and MES scores, we defined cutoffs separating the two distributions. Cells were then classified into ADRN_seq_ and MES_seq_ subgroups based on high versus low ADRN scores (**Fig. 1D**) and low versus high MES scores (**Fig. S1A**). These subgroups segregated into distinct regions in PCA space, with ADRN score-based subgrouping showing clearer separation. The relative abundance of these subgroups varied substantially across cell lines (**Fig. 1E**). Specifically, ADRN score–based subgrouping demonstrated greater heterogeneity and clearer separation, for example, in BE2C, IMR5, and SK-N-FI, suggesting that the ADRN score is a more effective criterion for clustering. Notably, average ADRN and MES scores across cell lines were inversely correlated (R = −0.674, P < 0.05; **Fig. 1F; Supplementary Data 2**).

We next assessed the fraction of ADRN_seq_ and MES_seq_ defined by ADRN scores (**Fig. 1G**) and MES scores (**Fig. S1B**) for each cell line. The fraction of ADRN_seq_ cells defined by the two scores was strongly correlated (R = 0.913, P < 0.001; **Fig. S1C; Supplementary Data 2**). Furthermore, the fraction of ADRN_seq_ showed a significantly positive correlation with average ADRN score derived from scRNA-seq (R = 0.964, P<0.001, **Fig. 1H**) and ADRN scores calculated from bulk RNA sequencing (RNA-seq) using ssGSEA (R = 0.714, P < 0.05, **Fig. S1D; Supplementary Data 2**). Although not statistically significant, the average ADRN scores from scRNA-seq showed positive correlation with the ADRN score from bulk RNA-seq (**Fig S1E**). Together, these results identify robust ADRN and MES transcriptional states at single-cell resolution in NB cell lines, providing a definitive “ground truth” for subsequent morphology-based classification.

### Cell morphology reflects underlying NB transcriptional states

To confirm the differences in molecular features between ADRN_seq_ and MES_seq_ subgroups (hereafter defined by ADRN score), we examined the expression of canonical lineage markers^12,13^. MES markers, including yes-associated protein 1 (*YAP1*) and vimentin (*VIM*), were preferentially expressed in MES_seq_ cells, whereas ADRN markers such as GATA-binding protein 3 (*GATA3*) and paired-like homeobox 2B (*PHOX2B*) were enriched in ADRN_seq_ cells (**Fig. 2A**). This pattern was consistently observed across NB cell lines, with expression levels correlating with the fraction of ADRN_seq_ cells in each line (**Fig. 2B**). To further investigate phenotypic features of MES_seq_ cells distinct from ADRN_seq_ cells, we performed Gene Ontology (GO) enrichment analysis on genes highly expressed in this subgroup (**Supplementary Data 3**) using Cytoscape and ClueGo ^20,21^. Enriched biological processes included actin filament organization, cell adhesion, epithelial morphology, and cell spreading (**Fig. 2C**), suggesting that unique MES_seq_ characteristics are strongly coupled to cell morphology.

**Figure 2.**
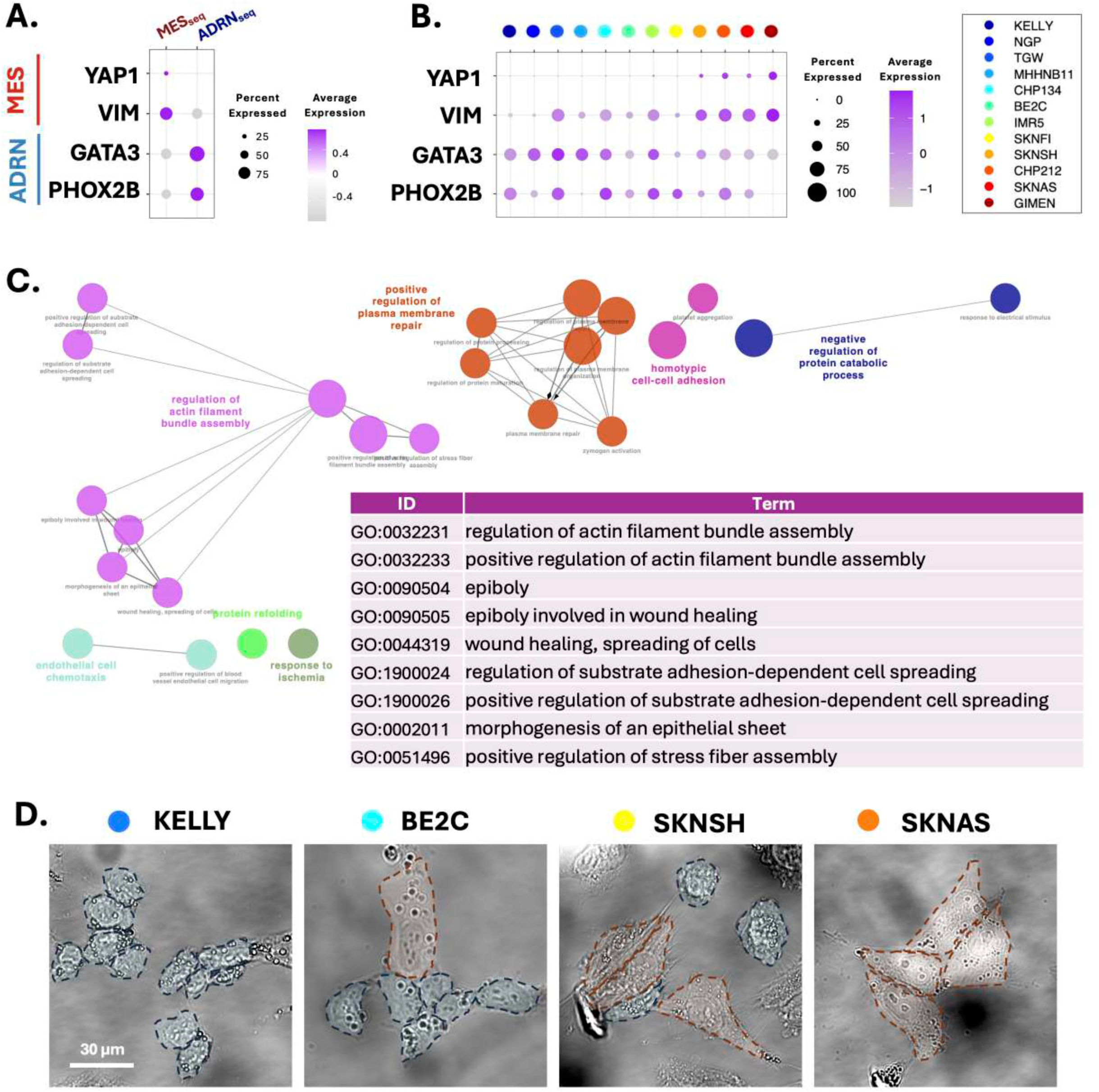
Higher expression of cell morphology-associated genes in MES_seq_ subgroup. **(A)** Dot plot showing the expression of MES markers (YAP1 and VIM) and ADRN markers (GATA3 and PHOX2B) in MES_seq_ and ADRN_seq_ subgroups in Fig. 1G. **(B)** Dot plot showing the expression of MES markers (YAP1 and VIM) and ADRN markers (GATA3 and PHOX2B) across individual cell line ordered by the fraction of ADRN_seq_ **(C)** Gene Ontology (GO) analysis of highly expressed genes in MES_seq_ using Cytoscape. **(D)** Representative morphologies of NB cell lines. Red dashed lines indicate MES-like cells, whereas blue dashed lines indicate ADRN-like cells.

Given this association, we next examined whether cell morphology could serve as a reliable readout of NB cell states. Cell morphology is an easily observable phenotype that integrates molecular and structural features linked to cancer progression. Notably, MES-associated molecular features are highly complex (**Fig. S2**), highlighting the need for more accessible phenotypic readouts. Consistent with this, NB cell lines displayed two distinct morphologies under standard culture conditions: a rounded, clustered phenotype (ADRN-like) and an elongated, spread phenotype (MES-like) (**Fig. 2D**).

To determine whether morphology can define discrete cell states, we constructed a multidimensional feature space (“morphological fingerprints”) capturing cell morphology (**Supplementary Data 4**). We reasoned that if morphology encodes cell identity, cells should organize into separable regions within this feature space. These features included cell spreading area, major axis, minor axis, elongation factor, perimeter, and circularity. After normalization, dimensionality reduction was performed using PCA, followed by unsupervised clustering using spectral clustering (**Fig. 3A**). Spectral clustering is uniquely suited for capturing complex, non-linear relationships in high-dimensional morphological data ^22,23^, making it advantageous over k-means clustering, which we used for ADRN/MES score-based classification, and which relies on simple distance to cluster centroids.

**Figure 3.**
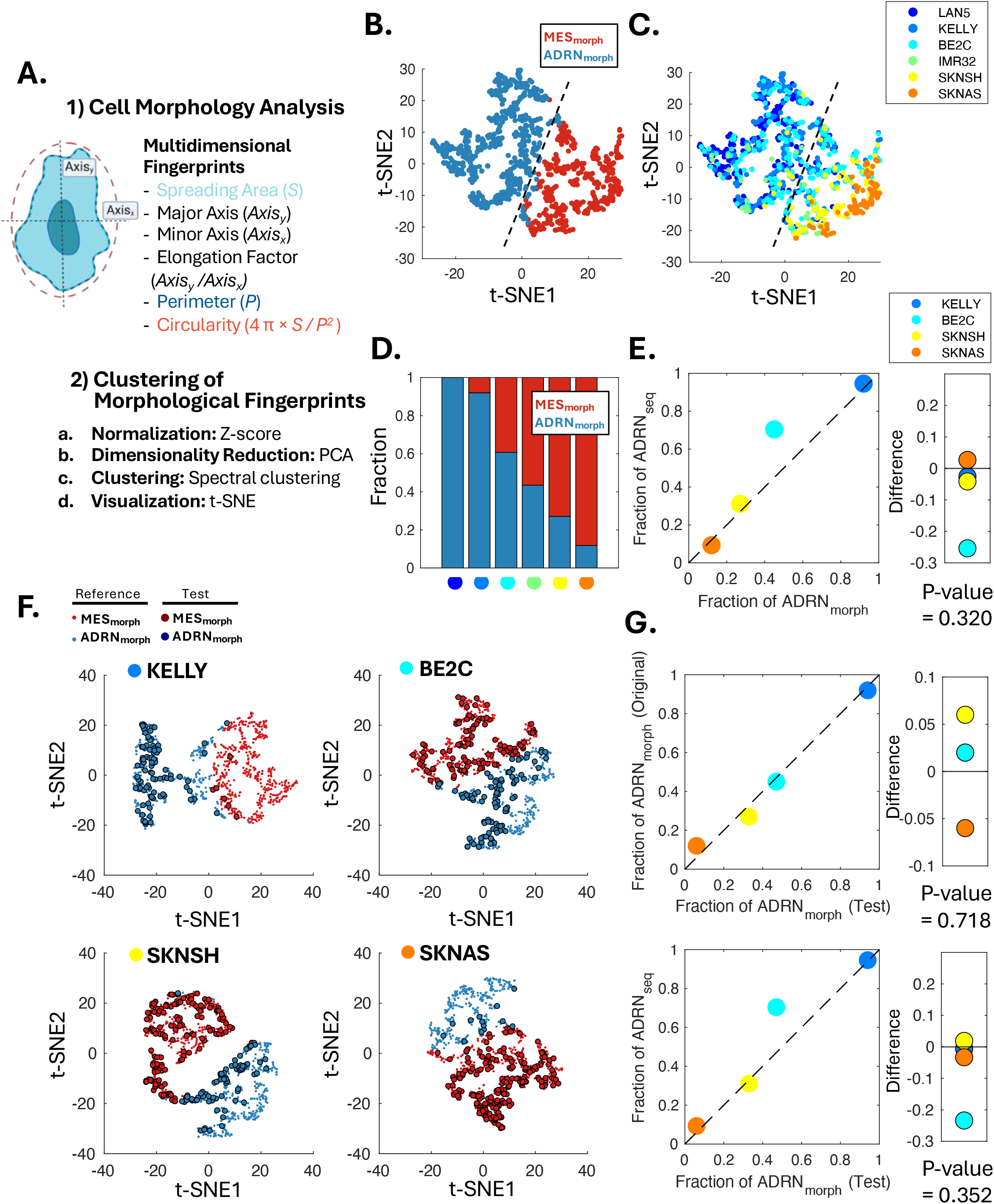
Morphology-based clustering identifies ADRN- and MES-like cell states in neuroblastoma cell lines. **(A)** Schematic overview of the cell morphology analysis workflow. Multidimensional morphological features (“morphological fingerprints”) were extracted and performed via unsupervised clustering (spectral clustering) after normalization and dimensionality reduction. **(B)** t-SNE plot showing two clusters identified based on morphological fingerprints, corresponding to ADRN_morph_ and MES_morph_ cells. **(C)** t-SNE plot showing the distribution of cells from individual NB cell lines across morphology-based clusters. Colors indicate different cell lines. **(D)** Fraction of ADRN_morph_ and MES_morph_ cells in each cell line. Colored dots indicate corresponding cell lines in **Panel C**. **(E)** Scatter plot comparing the fraction of ADRN_morph_ cells with the fraction of ADRN_seq_ cells derived from scRNA-seq analysis across cell lines that have scRNA-seq results. The difference between the two measurements is not statistically significant evaluated by one-sided t-test. **(F)** Cross validation of each test cell line. Test cells (large dots) from excluded test lines are projected onto the reference state space (small dots) to evaluate model generalizability. **(G)** Statistical consistency of cross validation. Scatter and difference plots compare validation results with the original model (top, P-value = 0.718) and scRNA-seq ground truth (bottom, P-value = 0.352), which are not statistically significant evaluated by one-sided t-test.

Indeed, projection of cells into this space revealed two distinct clusters, corresponding to elongated, spread morphologies (MES-like) and compact, rounded morphologies (ADRN-like) (**Fig. 3B**). We therefore define this morphology-derived embedding as a state space, in which cell identity can be inferred without transcriptomic measurements. The cluster of cells characterized by more elongated and spread morphologies was defined as MES_morph_, whereas the other cluster exhibiting rounded and compact morphologies was defined as ADRN_morph_. We then quantified the fraction of ADRN_morph_ cells in each cell line and compared it with the fraction of ADRN_seq_ cells obtained from scRNA-seq analysis (**Fig. 3C and D**). Notably, the two measurements were well aligned, with no statistically significant difference between the morphology-based and transcriptomics-based fractions (**Fig. 3E**).

To ensure that our morphology-based inference framework was not overfitted to the specific characteristics of the cell lines used for initial clustering, we performed a cross validation. For each iteration, we constructed a reference morphological state space excluding one testing cell line and then projected the “unseen” test cells from the excluded line onto this reference space. Visual inspection of the t-SNE projections confirmed that test cells from diverse genetic backgrounds and morphology (KELLY, BE2C, SK-N-SH, and SK-N-AS) were accurately mapped onto the pre-defined morphological topology, aligning closely with the ADRN_morph_ and MES_morph_ regions established by the reference lines (**Fig. 3F**).

Statistical analysis further confirmed the robustness of this approach (**Fig. 3G**). The ADRN_morph_ fractions predicted by the cross validation were highly consistent with those derived from the original inclusive model (e.g., 0.92 vs. 0.94 for KELLY; 0.12 vs. 0.06 for SK-N-AS), with a paired t-test showing no significant difference between the two (P = 0.718). Crucially, the cross validateion-derived morphological fractions also remained statistically indistinguishable from the scRNA-seq ground truth (P = 0.352). These findings demonstrate that our framework captures universal phenotypic features of NB cell states rather than cell-line-specific artifacts, ensuring its generalizability to independent datasets.

We next evaluated whether morphology-based clusters exhibit distinct molecular features associated with ADRN and MES states. To assess these features at the single-cell level, NB cells were immunofluorescently stained for GATA3 (an ADRN marker) and YAP (an MES marker). GATA3 intensity and YAP activity were quantified in individual cells, with YAP activity defined as the nuclear-to-cytoplasmic ratio of YAP intensity (**Fig. 4A**).

**Figure 4.**
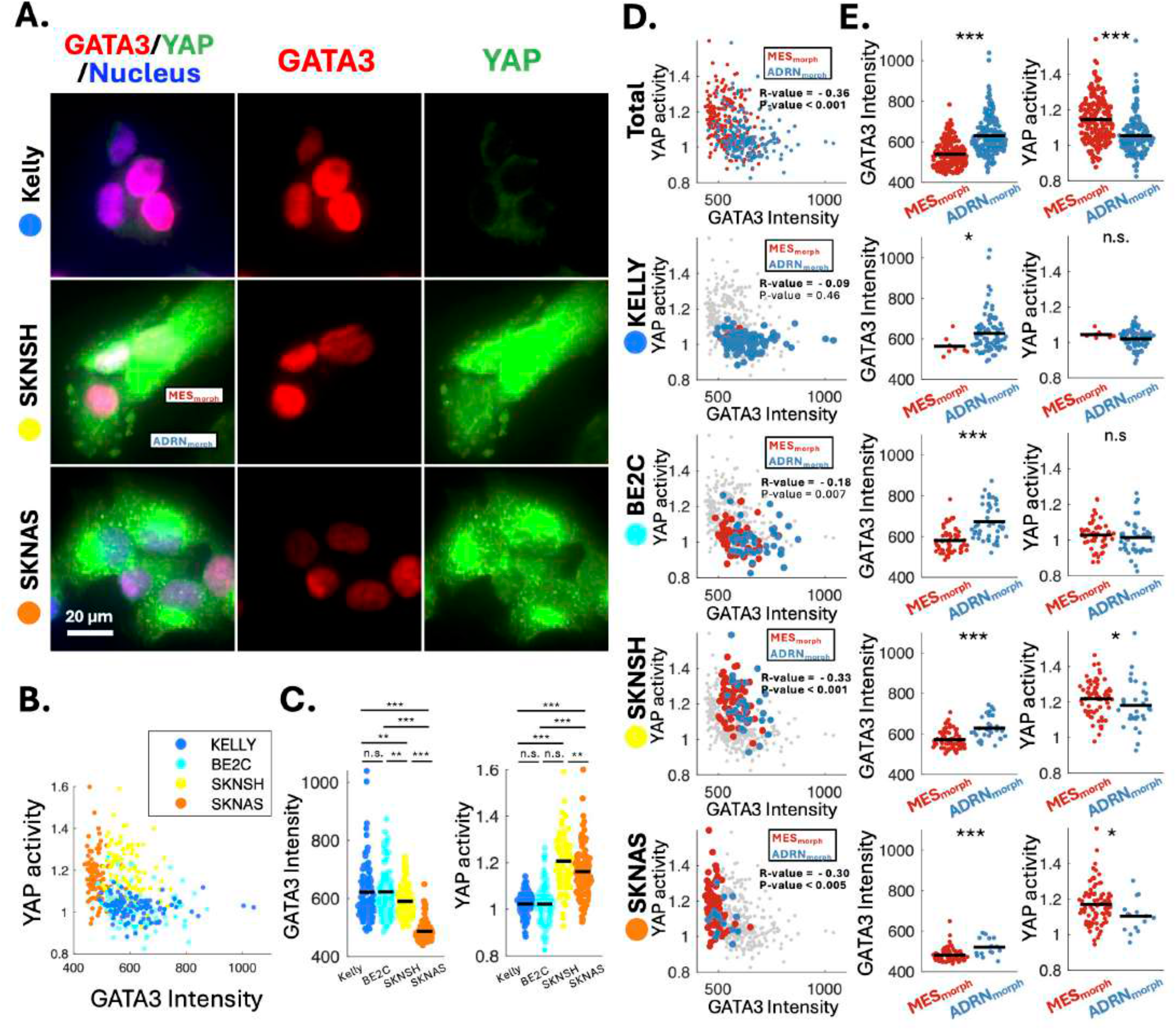
Molecular differences between ADRN_morph_ and MES_morph_ cells. **(A)** Immunofluorescence staining of NB cells for GATA3 (red; ADRN marker) and YAP (green; MES marker), with nuclei shown in blue (Hoechst). **(B)** Scatter plot showing the relationship between GATA3 intensity and YAP activity. YAP activity is defined as the ratio of nuclear to cytoplasmic YAP intensity. **(C)** Quantification of GATA3 intensity (left) and YAP activity (right) across NB cell lines, ordered by the fraction of ADRN_morph_ cells. Statistical significance was assessed using one-way ANOVA.*P < 0.05, **P < 0.01, ***P < 0.005, ****P < 0.001; n.s., not significant. **(D)** Scatter plots of GATA3 intensity versus YAP activity in MES_morph_ and ADRN_morph_ cells, shown for the combined dataset and for each individual cell line. Correlation coefficients (R) and P values were calculated using Pearson’s correlation analysis. **(E)** Quantification of GATA3 intensity (left) and YAP activity (right) in MES_morph_ and ADRN_morph_ cells for the combined dataset and for each cell line. Statistical significance was assessed using a two-sided Student’s *t*-test. *P < 0.05, **P < 0.01, ***P < 0.005, ****P < 0.001; n.s., not significant.

Consistent with the opposing relationship between ADRN and MES transcriptional programs, GATA3 intensity and YAP activity were inversely correlated across cells, consistent with the opposing relationship between ADRN and MES transcriptional programs (R = −0.36, P < 0.001; **Fig. 4B and D, Supplementary Data 5**), consistent with the opposing ADRN and MES transcriptional programs. Cell lines with a higher fraction of MES_morph_ cells exhibited lower GATA3 intensity and higher YAP activity (**Fig. 4C**). Furthermore, ADRN_morph_ cells within each cell line displayed higher GATA3 intensity, indicative of stronger ADRN molecular features, compared to MES_morph_ cells. Although not statistically significant in KELLY and BE2C, YAP activity tended to be higher in MES_morph_ cells than in ADRN_morph_ cells (**Fig. 4D**).

To further validate these observations, we analyzed SK-N-SH cells, which exhibit substantial NB heterogeneity, using an additional pair of MES and ADRN markers (VIM and PHOX2B). Consistently, MES_morph_ cells showed higher VIM expression and lower PHOX2B intensity compared to ADRN_morph_ cells (**Fig. S3A and B; Supplementary Data 6**). To assess phenotypic differences between MES_morph_ and ADRN_morph_ cells, we examined cell migration using live-cell imaging. We projected the morphologies of SK-N-SH cells onto the reference morphological space defined in **Fig. 3**, assigning that ADRN_morph_ and MES_morph_ clusters (**Fig. S3C**). We then confirmed that MES_morph_ cells exhibited enhanced cell migration and pronounced displacement (**Fig. S3D and E, Supplementary Data 7**), consistent with MES-like phenotypic characteristics. Together, these results demonstrate that machine learning–based classification using morphological fingerprints reliably captures distinct MES and ADRN states in NB cells.

### ROCK signaling drives directional trajectories in morphology-defined state space

GO analysis revealed that genes highly expressed in MES_seq_ cells are significantly associated with Rho-associated kinase (ROCK) (**Fig. 5A**). ROCK is a well-established regulator of actin dynamics and cell morphology ^24,25^, and these findings are consistent with prior studies suggesting that the interaction of NB cells with extracellular matrix–mediated by ROCK contributes to NB cell state plasticity ^26^. We further confirmed that ROCK inhibition by Y27632 shifted transcriptional profiles away from an MES-like state, as evidenced by a reduction in MES scores (**Fig. S4A**).

**Figure 5.**
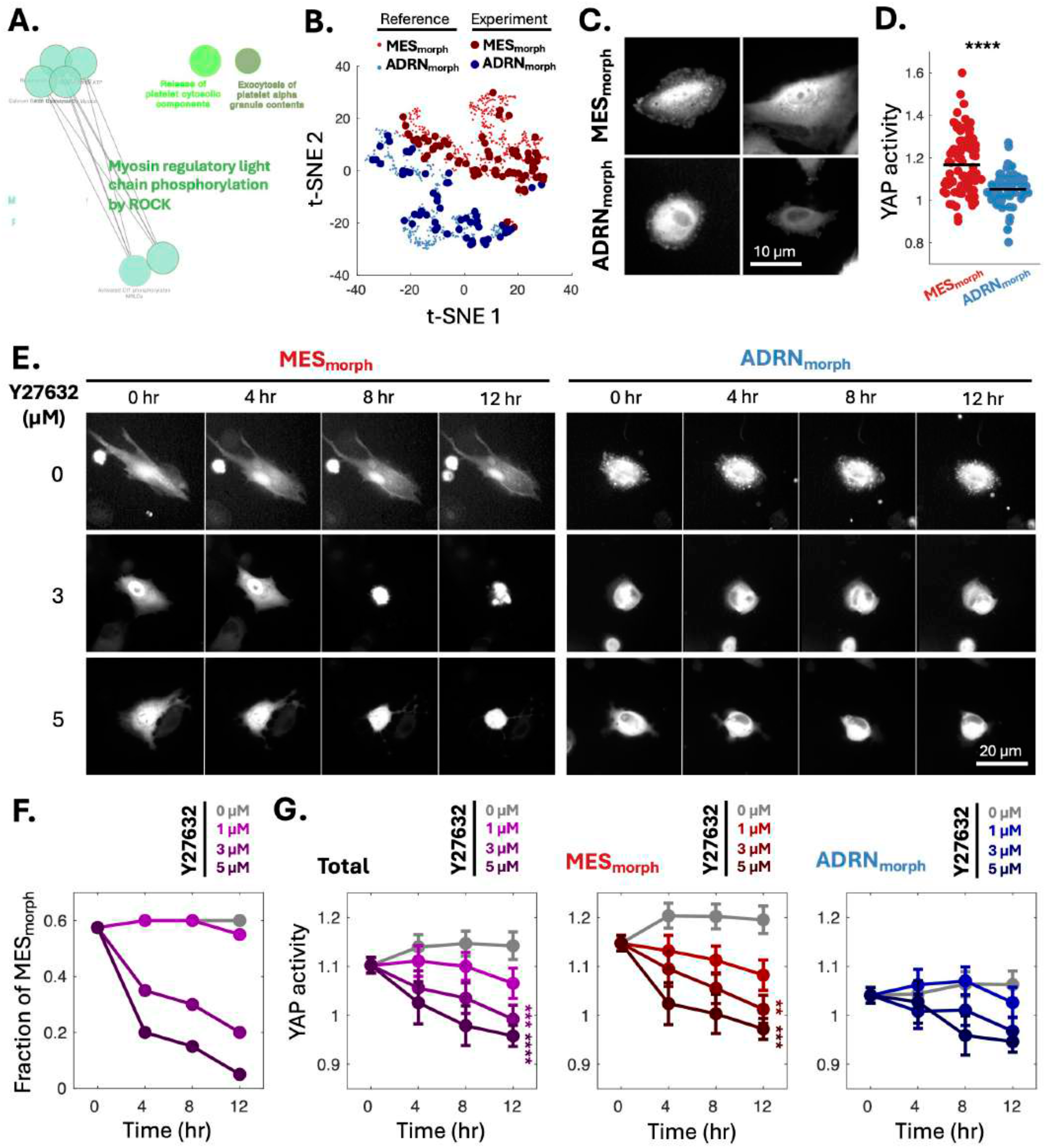
ROCK inhibition promotes transition of NB cells toward a ADRN_morph_ state. **(A)** GO analysis highlighting enrichment of ROCK-mediated myosin regulatory light chain phosphorylation among genes highly expressed in MES_seq_ cells, identified using Cytoscape. **(B)** t-SNE plot showing morphology-based clusters of reference cells (Fig. 3) and SK-N-SH cells expressing a YAP live-cell reporter (Experiment). **(C)** Representative live-cell images of YAP activity in MES_morph_ (top) and ADRN_morph_ (bottom) SK-N-SH cells. **(D)** Quantification of YAP activity, defined as the nuclear-to-cytoplasmic ratio of YAP intensity, in ADRN_morph_ and MES_morph_ SK-N-SH cells. Lines indicate group means. Statistical significance was assessed using a two-sided Student’s *t* test, ****P < 0.001. **(E)** Representative live-cell images showing YAP activity over 12 hours following treatment with Y27632 (a ROCK inhibitor) in MES_morph_ and ADRN_morph_ cells. **(F)** Time-resolved transitions across morphology-defined state space upon ROCK inhibition. Changes in the fraction of MES_morph_ cells at different concentrations of Y27632 over 12 hours. **(G)** Time-course changes in YAP activity in total cells, MES_morph_ cells, and ADRN_morph_ cells at different concentrations of Y27632. Statistical significance at 12 hours was assessed by one-way ANOVA, **P < 0.01, ***P < 0.005.

Based on these observations, we hypothesized that ROCK signaling functionally modulates transitions between NB cell states. To test this, we used SK-N-SH cells expressing a YAP live-cell reporter (EGFP-YAP), enabling real-time monitoring of signaling dynamics during state transitions. Projection of these cells onto the reference morphology-defined state space confirmed that ADRN_morph_ and MES_morph_ clusters were robustly preserved in the experimental dataset (**Fig. 5B**). YAP has been served as a MES marker and demonstrated that its activation drives the transition of ADRN to MES states ^26,27^. Consistent with the link between morphology and signaling, YAP activity differed between MES_morph_ and ADRN_morph_ cells (**Fig. 5C and D**), supporting its use as a functional readout of cell state.

We next treated cells with the ROCK inhibitor Y27632 and monitored morphology and YAP activity over time. ROCK inhibition induced dose- and time-dependent transitions from MES_morph_ to ADRN_morph_ states, following a continuous and directional trajectory within the morphology-defined state space (**Fig. 5E, 5F; Fig. S4B and S4C; Supplementary Data 8**). At the single-cell level, time-resolved tracking revealed coherent trajectories of individual cells progressing toward the ADRN_morph_ region, supporting the existence of a directional axis within this space (**Fig. S4B**). Correspondingly, the fraction of MES_morph_ cells decreased over time with increasing concentrations of Y27632, while ADRN_morph_ cells increased (**Fig. 5F; Fig. S4C**), indicating a global shift in population-level state distribution. These transitions were accompanied by progressive reductions in YAP activity (**Fig. 5G; Fig. S4D**), consistent with coordinated changes in underlying signaling programs. Specifically, MES_morph_ cells at the initial time points showed more frequent transitions toward ADRN_morph_ states, accompanied by significant reductions in YAP activity **(Fig. 5F; Fig. S4C and S4D)**, suggesting that ROCK inhibition has a more pronounced effect on MES states than on ADRN states.

Importantly, these transitions were captured at single-cell resolution using morphological features alone, without reliance on transcriptomic measurements. Together, these results demonstrate that perturbation of ROCK signaling drives continuous and directional movement within a morphology-defined state space, establishing a mechanistically grounded trajectory for cell state transitions. This trajectory provides a reference framework for comparing how distinct regulatory perturbations reshape cell identity within a shared phenotypic landscape.

### Epigenetic perturbation converges onto a shared morphology-defined state trajectory^28^

To examine how epigenetic regulation shapes NB cell identity, we analyzed H3K27me3 deposition across ADRN and MES signature loci. Our analysis of chromatin immunoprecipitation sequencing (ChIP-seq) data publicly accessible (GSE138314) revealed that ADRN-associated loci are differentially marked across NB cell lines, with H3K27me3 levels inversely correlating with ADRN transcriptional scores. In contrast, H3K27me3 enrichment at MES-associated loci showed no significant correlation with ADRN scores, suggesting that the Polycomb-mediated repressive landscape preferentially defines the ADRN program rather than the MES state (**Fig. 6A, B; Fig. S5A and S5B; Supplementary Data 9**).

**Figure 6.**
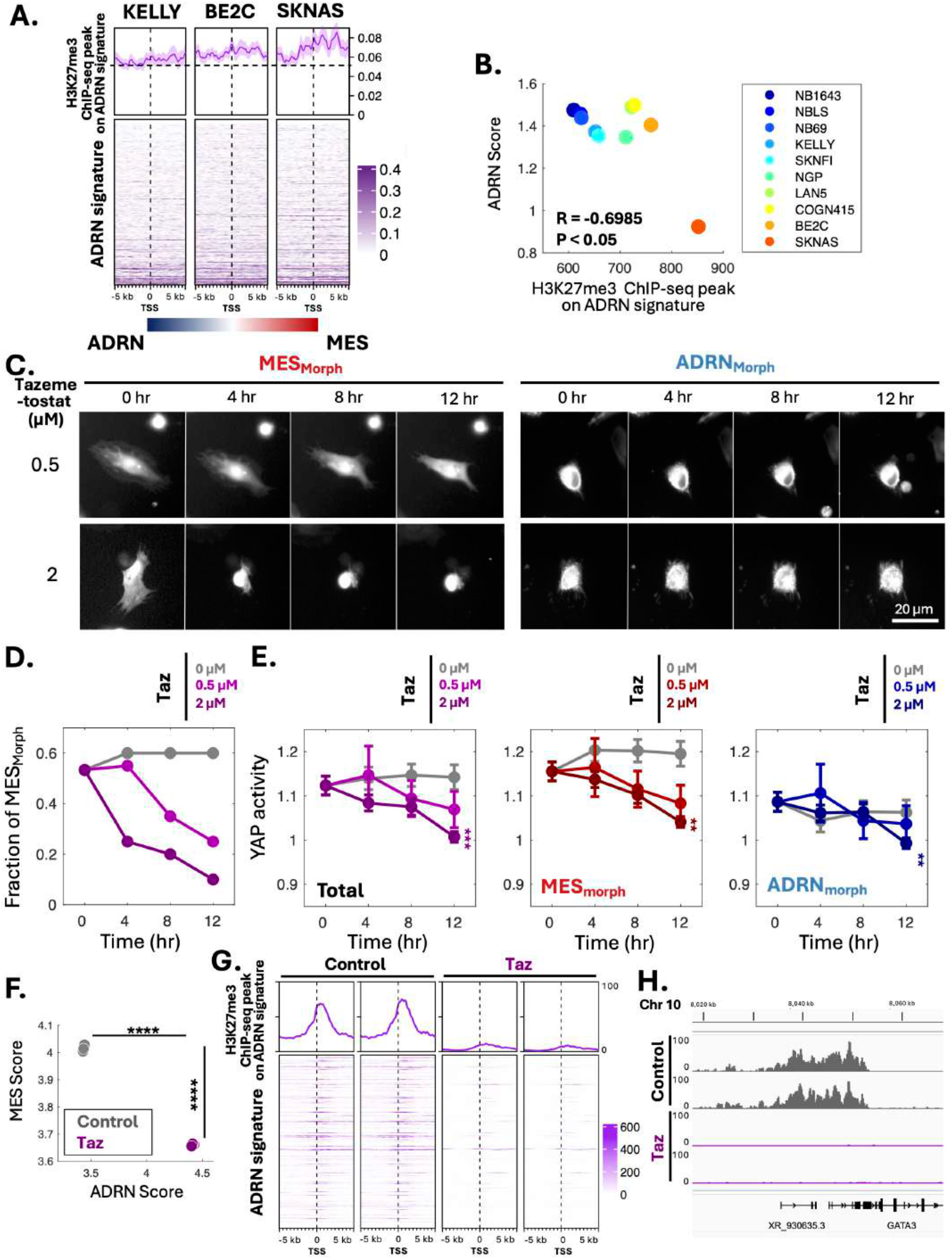
EZH2 inhibition promotes transition of NB cells toward an ADRN_morph_ state. **(A)** H3K27me3 chromatin immunoprecipitation sequencing (ChIP-seq) signal (top) and heatmaps (bottom) at ADRN signature loci in NB cell lines (KELLY, BE2C, and SK-N-AS) with available ChIP-seq and morphology data. **(B)** Scatter plot showing the relationship between H3K27me3 ChIP-seq peak intensity at ADRN signature regions and ADRN scores derived from bulk RNA-seq across NB cell lines. R and P values were calculated using Pearson’s correlation coefficient. **(C)** Representative live-cell images showing YAP activity over 12 hours following treatment with Tazemetostat in MES_morph_ and ADRN_morph_ cells. **(D)** Time-resolved transitions across morphology-defined state space upon EZH2 inhibition. Changes in the fraction of MES_morph_ cells at different concentrations of Tazemetostat over 12 hours. **(E)** Time-course changes in YAP activity in total cells, MES_morph_ cells, and ADRN_morph_ cells at different concentrations of Tazemetostat. Statistical significance at 12 hours was assessed using one-way ANOVA, **P < 0.01 and ***P < 0.005. **(F)** Scatter plot of ADRN and MES scores in SK-N-AS cells with and without Tazemetostat treatment (EZH2 inhibitor). Statistical significance was assessed using a two-sided Student’s t-test, ***P < 0.005. **(G)** H3K27me3 ChIP-seq signal (top) and heatmaps (bottom) at ADRN signature loci in SK-N-AS cells upon Tazemetostat treatment. **(H)** H3K27me3 ChIP-seq signal profile on GATA3 in SK-N-AS cells with and without Tazemetostat treatment.

Previous studies have shown that the pharmacological inhibition of EZH2, a component of Polycomb repressive complex 2, suppresses the expression of MES-associated molecules, including GD2. Given this, we assessed whether EZH2 inhibition using tazemetostat could drive state transitions in NB cells. Similar to the effects of ROCK inhibition, EZH2 inhibition induced dose- and time-dependent transitions from MES_morph_ to ADRN_morph_ states, following a continuous trajectory within the morphology-defined state space (**Fig. 6C, 6D; Fig. S5C and S5D; Supplementary Data 10**). These transitions were accompanied by a progressive reduction in YAP activity, consistent with a shift in underlying molecular features toward an ADRN-like state (**Fig. 6E; Fig. S5E**). Notably, EZH2 inhibition exerted a more pronounced effect on MES states, as mirrored by both morphological mapping and YAP activity, compared to ADRN states (**Fig. 6D; Fig. S5D and S5E**).

Importantly, these trajectories closely recapitulated those observed under ROCK inhibition, despite targeting a mechanistically distinct regulatory layer (**Fig. 5; Fig. S4**). This convergence was further validated at the transcriptomic level, where EZH2 inhibition significantly increased ADRN scores while decreasing MES scores (**Fig. 6F**). Furthermore, H3K27me3 occupancy at ADRN signature loci, e.g. GATA3, was suppressed upon EZH2 inhibition, whereas changes at MES loci were less dramatic (**Fig. 6G and H; Fig. S5F**). Together, these results demonstrate that our morphology-based workflow accurately predicts epigenetic reprogramming toward an ADRN state. These findings suggest that morphology defines a low-dimensional state space onto which distinct regulatory mechanisms, including signaling and epigenetic programs, collapse, providing a unified and scalable representation of cancer cell identity.

## Discussion

In this study, we establish a morphology-based machine learning framework that enables inference of cancer cell states directly from imaging data, without reliance on transcriptomic profiling. We show that cell morphology defines a low-dimensional latent space that quantitatively encodes cellular identity and its dynamics, with morphology-defined states closely aligning with transcriptional classifications at single-cell resolution. Specifically, we show that NB cellular morphology serves as a scalable and quantitative proxy to distinguish ADRN and MES states at single-cell resolution.

Furthermore, we demonstrate that this framework captures dynamic cell state transitions across mechanistically distinct perturbations. Both ROCK inhibition and EZH2 inhibition induced time- and dose-dependent shifts in morphology-defined states, accompanied by coordinated changes in YAP activity, a molecular readout of NB cell identity. These perturbations converged on consistent morphological readouts, suggesting that morphology integrates downstream effects of diverse regulatory programs. Taken together, these findings support a model in which morphology functions as an integrated readout of multi-layer regulatory processes, including signaling and epigenetic states, effectively compressing high-dimensional molecular information into an observable phenotypic space.

Our analysis further suggests that the ADRN regulatory landscape more clearly defines NB cell states. ADRN scores exhibit a distinct bimodal distribution and more robust separation of subpopulations compared to MES scores, which display broader and less discrete patterns. In addition, epigenetic features are more clearly resolved within ADRN signatures than within MES signatures. These findings support a model in which the ADRN program represents a more stable and well-defined transcriptional state maintained by core regulatory circuitries, whereas MES identity reflects a more plastic continuum of transitional phenotypes ^29,30^. Consistent with this asymmetry, ADRN-defining super-enhancers, including those associated with PHOX2B and HAND1/2, are well characterized, while MES-associated regulators (e.g., AP-1) remain less clearly defined. In line with this, morphology-based classification more closely aligns with ADRN-defined states, supporting the use of ADRN-centric criteria for defining NB cell identity.

Unlike transcriptomic approaches, which are destructive and limited to static snapshots, morphology provides a real-time, non-invasive, and scalable readout of cellular state. This enables continuous tracking of cell state transitions at single-cell resolution and offers a practical strategy for studying cellular plasticity and therapeutic response dynamics. Although demonstrated here in NB, this framework is not restricted to a specific system. Rather, morphology may serve as a generalizable coordinate system for cellular state, particularly for transitions associated with cytoskeletal remodeling, such as epithelial-mesenchymal transition ^31-33^. Integration with imaging-based and multi-omic approaches may further extend its utility for large-scale analysis of cell state dynamics.

Together, our findings establish cell morphology as a low-dimensional, experimentally accessible state space that encodes both cellular identity and its dynamic transitions, providing a scalable foundation for image-based analysis of cellular plasticity.

## Materials and Methods

### Single-Cell RNA Sequencing Data Preprocessing and Integration

We analyzed single-cell RNA sequencing data of neuroblastoma (NB) cell lines obtained from GSE229224. Bulk RNA sequencing data were acquired from GSE89413 (Cell lines), Chronopolous and Vemula et al ^26^ (ROCK inhibition), and GSE180515 (EZH2 inhibition). To profile transcriptional landscapes across multiple NB cell lines, raw sequencing matrices, including matrix, feature, and barcode files, were systematically processed using the Seurat R package. Individual sample datasets were imported using the ReadMtx function to construct discrete Seurat objects. To resolve potential cell barcode overlaps across distinct experiments, unique cell identifiers were appended using sample-specific prefixes during dataset merging. The merged Seurat object was subsequently evaluated for quality control metrics. To filter out low-quality cells, damaged cells, or empty droplets, cells were excluded if their extracted genes per cell fell outside the range of 200 to 7500, or if the percentage of mitochondrial transcripts exceeded 10%. This mitochondrial threshold, calculated using the PercentageFeatureSet function targeting human mitochondrial genes prefixed with “MT-”, was utilized to eliminate apoptotic cells.

### Normalization, Dimensional Reduction, and Clustering

Following quality filtering, the single-cell expression matrix was normalized using a global scaling method with a scale factor of 10,000 and log-transformed via the NormalizeData function. Highly variable features were identified using the variance-stabilizing transformation method to select the top 2,000 variable genes. To prevent highly expressed transcripts from dominating downstream dimensional reductions, expression levels of these variable genes were scaled and centered to a mean of 0 and a variance of 1 across all cells. Principal Component Analysis (PCA) was performed on the scaled highly variable gene space. To determine statistically significant principal components contributing to biological variation, a JackStraw permutation test was implemented using 100 replicates via the JackStraw and ScoreJackStraw functions. Empirical p-values for genes associated with PC1 and PC2 were extracted directly from the Seurat object’s JackStraw slot and filtered at a significance threshold of less than 0.05 to characterize driving axes of variation.

### Phenotypic Signature Scoring and Thresholding

To characterize adrenergic (ADRN) and mesenchymal (MES) phenotypes across single cells, raw count matrices were consolidated across layers using JoinLayers and subjected to single-sample Gene Set Enrichment Analysis (ssGSEA) using the GSVA R package. The reference gene sets of ADRN and MES signatures for ssGSEA are from van Groningen et al ^13^.

To define the thresholds in a data-driven manner, ADRN and MES scores were partitioned into low- and high-expressing cohorts using unsupervised one-dimensional k-means clustering (k = 2). The classification thresholds for each program were mathematically defined as the maximum signature score within their respective low-expressing clusters, classifying any cell exceeding these boundary values as phenotype-high.

### Differential Expression and Marker Analysis

To identify transcriptionally distinct subpopulations, differential gene expression analyses were executed under two separate stratification paradigms. For cluster-based differential expression, we discovered markers defining the two-cluster k-means partition utilizing an optimized run of the Wilcoxon Rank-Sum test via the FindAllMarkers function. This analysis was accelerated by downsampling to a maximum of 1,000 cells per identity cluster while utilizing zero thresholds for both log-fold change and detection percentage to ensure a comprehensive, un-truncated gene output. For phenotype-based differential expression, cells were stratified into high and low ADRN cohorts and differential expression between these two cohorts was evaluated using the FindMarkers function with a Wilcoxon Rank-Sum test, adjusting for technical covariates, specifically total RNA counts and mitochondrial percentage, within the latent variables framework.

Genes were restricted to a minimum detection rate of 10% in either group and a minimum natural log fold-change threshold of 0.25.

We identified highly expressed genes in MES_seq_ cells using thresholds of log_2_ fold change ≥ 2 and expression in at least 10% of cells. Using these genes, enrichment analysis was performed in Cytoscape (v3.10.4) with the ClueGO plugin (v2.5.10). Reference gene sets included Gene Ontology Biological Process (GO BP; **Fig. 2C**) and Reactome pathways (**Fig. 5D**), with a significance threshold of P < 0.05. In addition, MES gene signatures were analyzed using GO Biological Process with the same P-value cutoff (**Fig. S2**).

### Cell Lines and Cell Culture

The following NB cell lines were purchased from ATCC: SK-N-SH, SK-N-AS, SK-N-FI, BE(2)-C. The NB cell lines Kelly and LAN-5 were kind gifts from Dr. Yves DeClerck (Children’s Hospital Los Angeles, CA). NB cell lines were cultured in high glucose DMEM supplemented with 20% Fetal Bovine Serum and 1% penicillin/streptomycin (Gibco/ThermoFisher; Waltham, MA, USA). We authenticated cell lines via STR profiling (CHLA Stem Cell core).

### Immunofluorescent Staining

NB cells (30,000 – 50,000 per well) were seeded onto Nunc LabTek II 8-well chamber glass plates (ThermoFisher) pre-coated for 1 hr with 1 µg/mL collagen type III (Advanced BioMatrix; Carlsbad, CA, USA). Cells were incubated for 2 days to allow for adherence and normal growth. Cells were then fixed with 4% paraformaldehyde in PBS for 10 min, permeabilized with 0.1% Triton X-100 (Sigma-Aldrich; St. Louis, MO, USA) in PBS for 10 min, and blocked with 1% BSA-PBS (Sigma-Aldrich) for 45 min. Primary antibodies to GATA3, PHOX2B, YAP, or vimentin were diluted 1:200 in blocking solution and added to cells for 2 hrs at room temperature, followed by 3 PBS washes to remove unbound antibodies. We followed with 1 hr incubation at room temperature with 1:600 Alexa 488- and Alexa 594-labeled secondary antibodies (Invitrogen; Waltham, MA, USA) as well as 1:2000 Hoescht stain to visualize nuclei, diluted in blocking solution. Cells were washed 3 more times with PBS and 500 µL of fresh PBS was added to each well for imaging. Full details of the antibodies used are described in the table below.

Epifluorescence images were obtained using a Nikon Ti2 Eclipse inverted microscope equipped with a Plan Apochromat λ 40× objective lens (Nikon Instruments Inc.; Melville, NY, USA).

|  | Antibody | Host and Clonality | Vendor | Catalog Number | Dilution |
| --- | --- | --- | --- | --- | --- |
| Primary | GATA3 [EPR16651] | Rabbit monoclonal | abcam | ab199428 | 1:200 |
|  | PHOX2B (B-11) | Mouse monoclonal | Santa Cruz Biotechnology | SC-376997 | 1:200 |
|  | Vimentin (D21H3) | Rabbit monoclonal | Cell Signaling Technologies | 5741S | 1:200 |
|  | YAP (H-9) | Mouse monoclonal | Santa Cruz Biotechnology | SC-271134 | 1:200 |
| Secondary | Goat Anti-Mouse IgG, IgM H&L Alexa Fluor 488 | Goat polyclonal | Invitrogen | A10680 | 1:600 |
|  | Goat Anti-Rabbit IgG, IgM H&L Alexa Fluor 594 | Goat polyclonal | Invitrogen | A11012 | 1:600 |

### Morphological Fingerprints Extraction and Clustering Analysis

Morphological feature extraction and analysis were performed using custom MATLAB scripts (MathWorks, MA). For each cell, morphological fingerprints were extracted from immunofluorescence images using automated segmentation based on MATLAB functions *imbinarize* and *bwselect*, or by manual delineation of cell boundaries when necessary. All features were subsequently normalized across cells using z-score normalization implemented with the MATLAB function *zscore*, which standardizes each feature to zero mean and unit variance. Dimensionality reduction was performed using PCA via the MATLAB function *pca*, which projects the data onto orthogonal components capturing the maximum variance in the dataset. Unsupervised clustering was then conducted using spectral clustering implemented with the MATLAB function *spectralcluster*, with the number of clusters set to two.

For visualization, t-distributed stochastic neighbor embedding (t-SNE) was applied to the normalized feature matrix to project high-dimensional data into two dimensions. Clusters were annotated based on morphological characteristics: cells with elongated and spread morphologies were classified as MES_morph_, whereas cells with rounded and compact morphologies were classified as ADRN_morph_.

### Cell Migration Analysis

NB cells were seeded onto Nunc LabTek II 8-well chamber glass plates (ThermoFisher) pre-coated for 1 hr with 1 µg/mL collagen type III (Advanced BioMatrix) for live-cell imaging analysis. Time-lapse imaging was performed over 10 hrs using a Nikon Ti2 Eclipse inverted microscope equipped with a Plan Apochromat λ 40× objective lens and an Okolab UNO T-H-CO2 live-cell imaging chamber for environmental control (37C, humidity, 5% CO_2_; Okolab; Pozzuoli, NA, Italy). Images were acquired at 2 hr intervals. Cell tracking was performed manually for each time point using customized MATLAB scripts. The resulting time-series coordinates were used to calculate migration speed by averaging cell displacements across each time interval. Total displacement was defined as the Euclidean distance between the initial and final cell positions over the 10 hr imaging period.

### Drug Treatment

For assays with YAP and EZH2 inhibition, SK-N-SH cells were treated with varying concentrations of the ROCK1/2 inhibitor Y27632 (SCM075; Sigma-Aldrich) or the EZH2 inhibitor Tazemetostat (S7128; Selleck Chemicals; Houston, TX, USA) for 24 hrs. Live-cell imaging analysis was performed prior to and after adding the indicated drugs using a Nikon Ti2 Eclipse microscope with live-cell imaging chamber for environmental control.

### YAP Live-cell Sensor Transfection and Live Cell Imaging

SK-N-SH cells were transfected with the mEGFP-N1-YAP plasmid (kind gift from Shuguo Sun, Addgene plasmid #166457) using Lipofectamine 3000 (Invitrogen) according to manufacturer’s instructions. Briefly, the mEGFP-N1-YAP plasmid was amplified and isolated from an agar stab using the ZymoPURE II Plasmid Midiprep kit (Zymo Research; Irvine, CA, USA). SK-N-SH cells were seeded in 6 well plates (1 × 10^6^ per well) and transfected the following day with 2.5 µg of mEGFP-N1-YAP plasmid DNA per well in Opti-mem (Gibco/ThermoFisher). After 48 hrs, cells were collected and mEGFP-positive cells were isolated through fluorescent activated cell sorting using a BD FACSymphony 5-color cell sorter (Becton, Dickinson, and Company; Franklin Lakes, NJ, USA). Positive cells were then plated on Nunc LabTek II 8-well chamber glass plates (ThermoFisher) pre-coated for 1 hr with 1 µg/mL collagen type III (Advanced BioMatrix) for live-cell imaging analysis.

Time-lapse imaging was performed over 12 hrs using a Nikon Ti2 Eclipse microscope with a Plan Apochromat λ 20× objective lens with live-cell imaging chamber for environmental control. Images were acquired at 4 hr intervals. The GFP channel was used to monitor YAP intensity and cell morphology at the single-cell level. Nuclear and cytoplasmic YAP intensities were manually quantified at each time point using customized MATLAB scripts. YAP activity was defined as the ratio of nuclear to cytoplasmic fluorescence intensity.

### Chromatin Immunoprecipitation Sequencing Analysis

Publicly available H3K27me3 ChIP-seq datasets for NB cell lines and EZH2 inhibition were obtained from GSE138314 and GSE180515. The data sets were analyzed to assess the epigenetic regulation of ADRN and MES transcriptional programs. ChIP-seq signal intensities at genomic regions corresponding to ADRN and MES signature genes were quantified by aggregating normalized peak signals within ±5 kb of transcription start sites (TSS). For each cell line, average H3K27me3 signal intensities across signature loci were computed and compared with corresponding ADRN and MES transcriptional scores derived from bulk RNA-seq. Correlation analyses between H3K27me3 enrichment and transcriptional scores were performed using Pearson’s correlation coefficient. Statistical significance was assessed using two-sided tests. Data processing, statistical analysis, and visualization were conducted using R, utilizing standard packages

### Use of Artificial Intelligence Tools

An artificial intelligence (AI)-based language model (ChatGPT, OpenAI; GPT-5.3) was used to assist with code development in R and MATLAB, as well as for language editing and improving clarity of the manuscript. The AI tool was not used for data analysis, figure generation, or scientific interpretation. All results, analyses, and conclusions were generated and verified by the authors, who take full responsibility for the content.

## Supporting information

Manuscript.pdf

## Code Availability

All custom scripts used for data processing, analysis, and visualization were implemented in R and MATLAB. The code used in this study is available from the corresponding author upon reasonable request and will be made publicly available upon publication.

## Figure Legends

**Figure S1.**
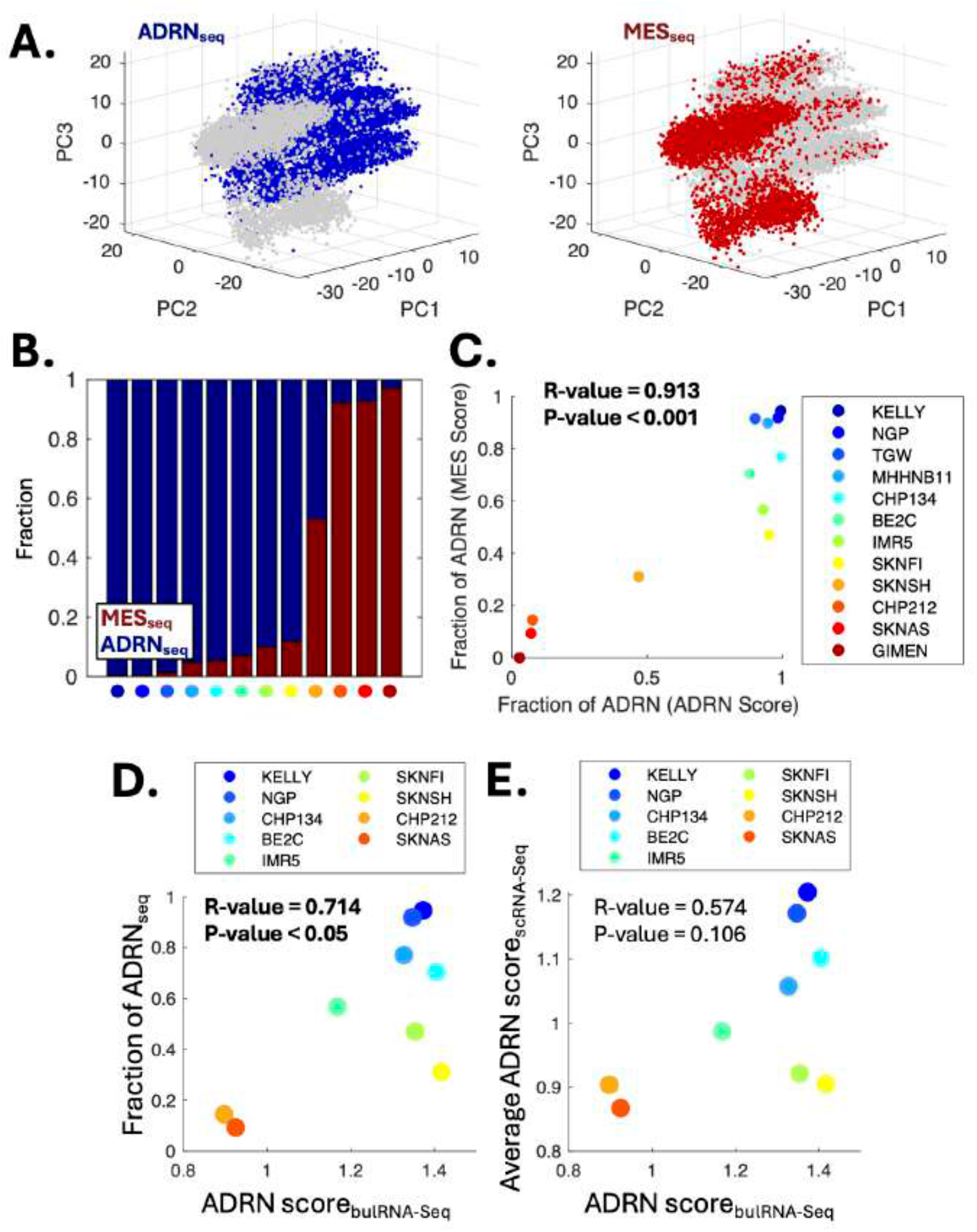
ADRN and MES subgroups defined by MES scores show consistency with bulk RNA sequencing results. **(A)** PCA plots highlighting an ADRN subgroup defined by low MES scores (ADRN_seq_, left) and an MES subgroup defined by high MES scores (MES_seq_, right) in the scRNA-seq analysis. **(B)** Fraction of ADRN_seq_ and MES_seq_ cells, defined based on MES scores, in each cell line. Each colored dot corresponds to a cell line shown in **Fig. 1F**. **(C)** Scatter plot showing the relationship between the fraction of ADRN_seq_ cells defined by high ADRN scores and those defined by low MES scores in each cell line. **(D)** Scatter plot showing the relationship between the fraction of ADRN_seq_ cells (defined by high ADRN scores) and ADRN scores derived from bulk RNA sequencing in each cell line. **(E)** Scatter plot showing the relationship between the average ADRN score from scRNA-seq and ADRN scores derived from bulk RNA sequencing in each cell line. In panels **(C), (D)**, and **(E)**, R and P values were calculated using Pearson’s correlation coefficient. Panels **(D)** and **(E)** use ADRN score–based subgrouping, consistent with Figure 1.

**Figure S2.**
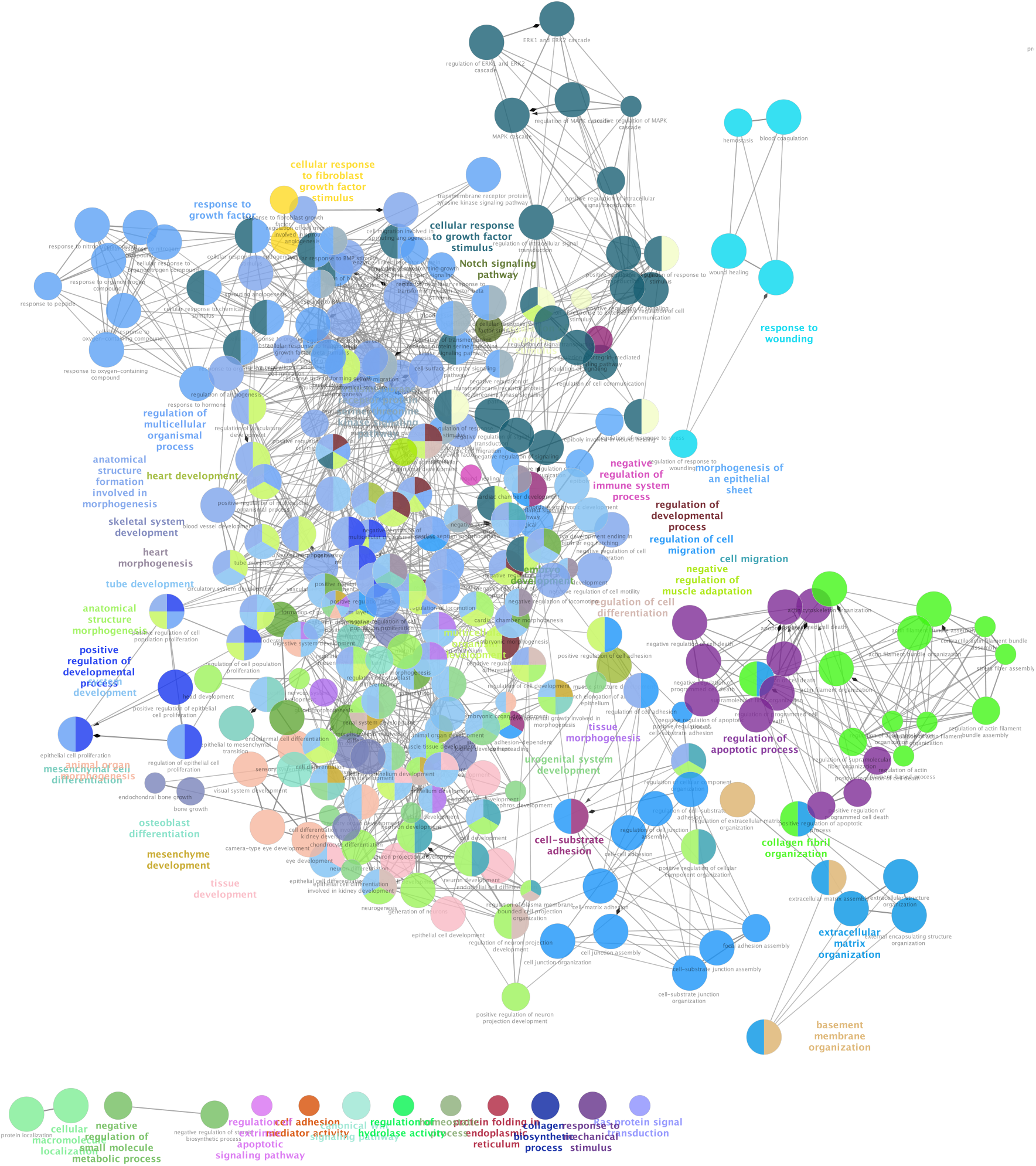
GO analysis of MES signatures using Cytoscape.

**Figure S3.**
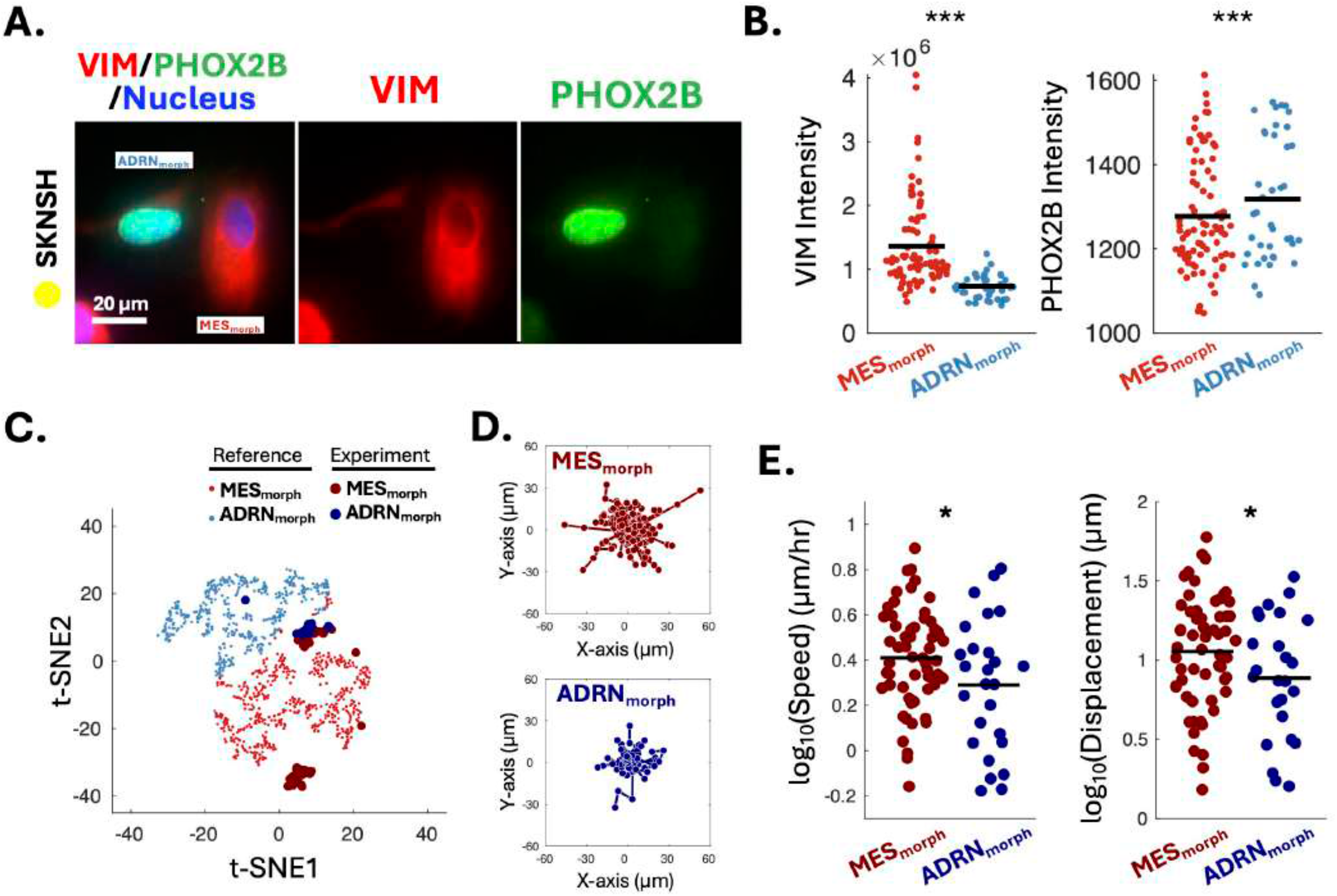
Distinct characteristics of ADRN_morph_ and MES_morph_ **cells in SK-N-SH**. **(A)** Immunofluorescence staining of SK-N-SH cells for VIM (red; MES marker) and PHOX2B (green; ADRN marker), with nuclei shown in blue (Hoechst). **(B)** Quantification of VIM (left) and PHOX2B intensity (right) in MES_morph_ and ADRN_morph_ cells. Statistical significance was assessed using a two-sided Student’s t-test. ***P < 0.005. **(C)** t-SNE plot showing morphology-based clusters of reference cells (Fig. 3) and SK-N-SH cells used for migration analysis (Experiment). **(D)** Migration trajectories of MES_morph_ and ADRN_morph_ cells over 10 hours from Experiment set in Panel C. **(E)** Quantification of migration speeds and displacements over 10 hours of MES_morph_ and ADRN_morph_ cells. Statistical significance was assessed using a two-sided Student’s *t*-test. *P < 0.05.

**Figure S4.**
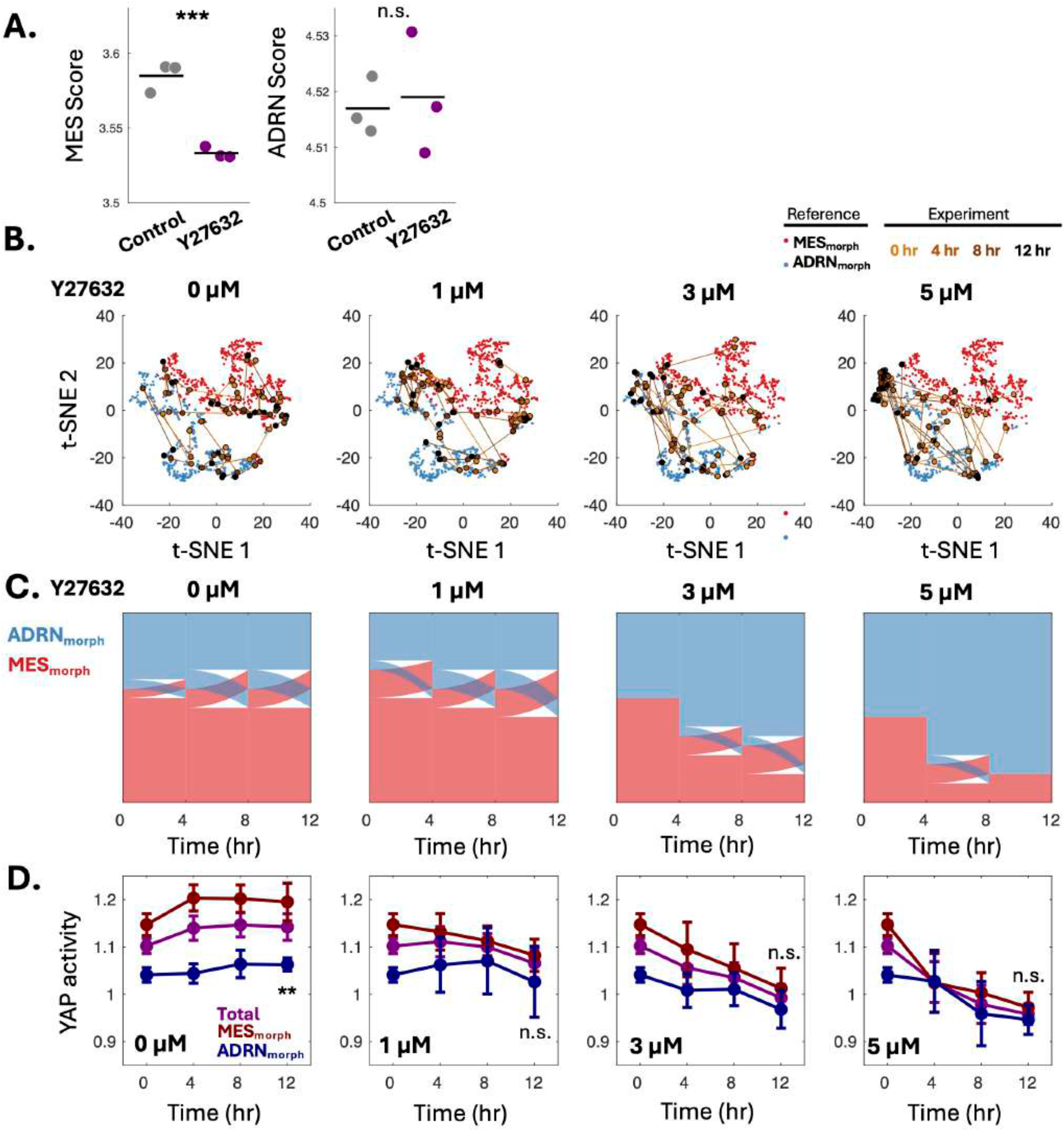
ROCK inhibition modulates morphology and reshapes NB states. **(A)** MES and ADRN scores following treatment with Y27632. Statistical significance was assessed using a two-sided Student’s t-test, ***P < 0.001 and n.s. = not significant. **(B)** t-SNE plots showing the distribution of cells across morphology-based clusters (ADRN_morph_ and MES_morph_) under increasing concentrations of Y27632 (0, 1, 3, and 5 μM) over time (0–12 h). Lines indicate cell trajectories across time points. Reference clusters from Fig. 3 are shown alongside experimental data. **(C)** Time-resolved changes in the fraction of ADRN_morph_ (blue) and MES_morph_ (red) cells at different concentrations of Y27632. **(D)** Time-course of YAP activity in total cells (purple), MES_morph_ cells (red), and ADRN_morph_ cells (blue) under increasing concentrations of Y27632. Statistical significance at 12 hours points was assessed by a two-sided Student’s t-test, **P-value = 0.01 and n.s.= not significant.

**Figure S5.**
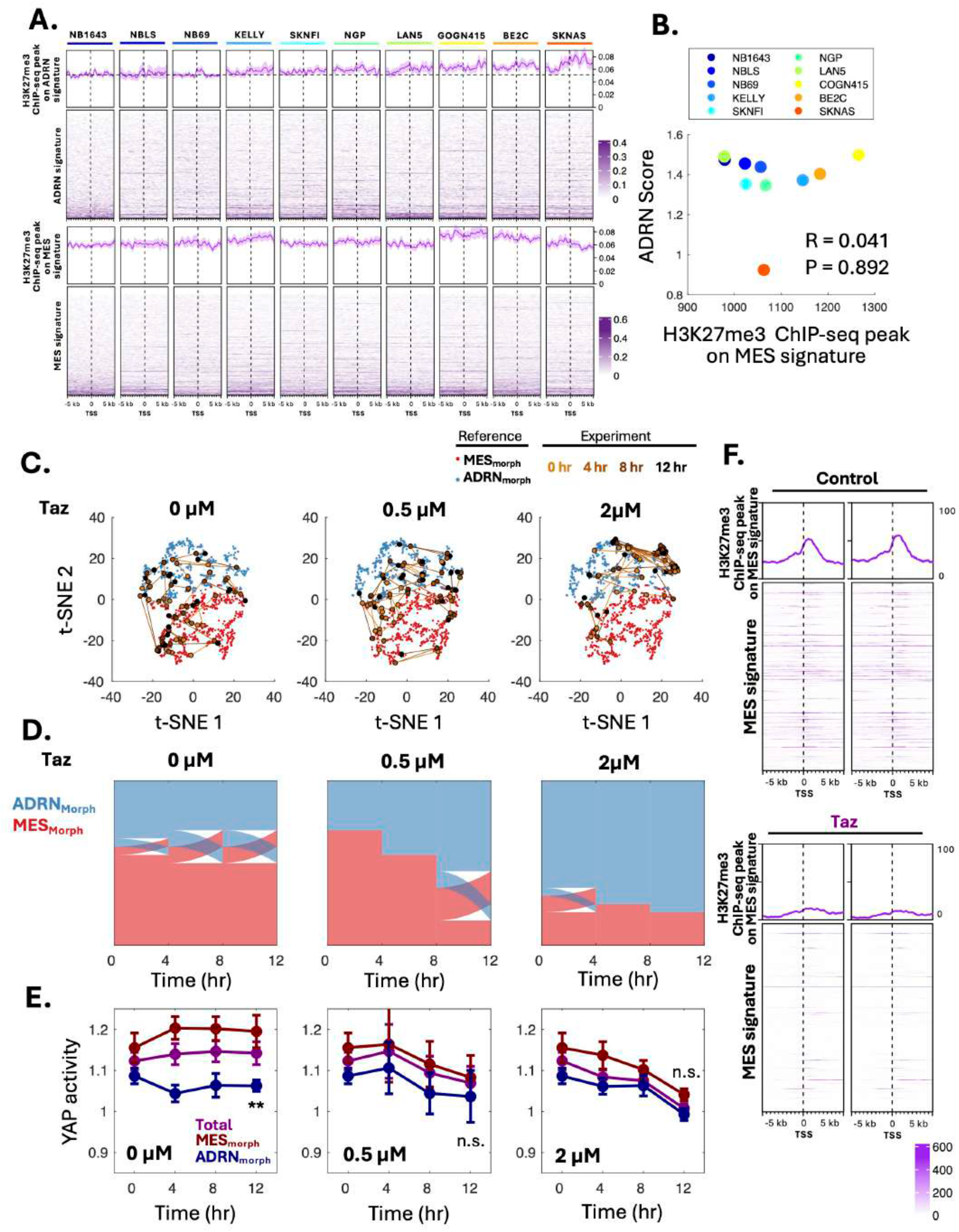
EZH2 inhibition modulates chromatin states at ADRN signatures and promotes ADRN-associated morphological and signaling features. **(A)** H3K27me3 ChIP-seq signal (top) and heatmaps (bottom) at ADRN (upper panels) and MES (lower panels) signature loci across multiple NB cell lines. **(B)** Scatter plot showing the relationship between H3K27me3 ChIP-seq peak intensity at MES signature regions and ADRN scores derived from bulk RNA-seq across NB cell lines. R and P values were calculated using Pearson’s correlation coefficient. **(C)** t-SNE plots showing morphology-based clusters under treatment with increasing concentrations of Tazemetostat (0, 0.5, and 2 μM) over time (0–12 h). Black lines indicate cell trajectories across time points. Reference clusters are shown alongside experimental data. **(D)** Time-resolved changes in the fraction of ADRN_morph_ (blue) and MES_morph_ (red) cells at different concentrations of Tazemetostat. **(E)** Time-course of YAP activity in total cells (purple), MES_morph_ cells (red), and ADRN_morph_ cells (blue) at different concentrations of Tazemetostat. Statistical significance at 12 hours points was assessed by a two-sided Student’s t-test, **P-value = 0.01 and n.s.= not significant. **(F)** H3K27me3 ChIP-seq signal (top) and heatmaps (bottom) at MES signature loci in SK-N-AS cells upon Tazemetostat treatment.

## Acknowledgements

This work was supported by the following funds: CHLA start-up fund, CHLA Core pilot grant, Margaret E. Early Medical Research Trust, National Cancer Institute under Grant P01 CA217959 (PI J. Maris) to J.P (PI).

## Author Contributions

V.Z. performed data curation, data analysis, writing, review, and editing. Y.Z. performed data analysis. J.P. performed conceptualization, supervision, funding acquisition, and data analysis, and wrote, re-viewed, and edited the final manuscript.

